# Potential geographic distribution and ecological niche of New World dobsonflies (Megaloptera, Corydalidae): the case of the Neartic-Neotropical transition zone

**DOI:** 10.1101/2022.01.01.474714

**Authors:** Hugo Alejandro Álvarez, Miguel A. Rivas-Soto

## Abstract

The dobsonflies (Megaloptera, Corydalidae) are an interesting group of insects, but among the New World dobsonflies, it is not known the effect of the Neartic-Neotropical transition zone on their biogeographic distribution. Here we studied at the species level, the records on the geographic range of the dobsonflies that occurred in and near the transition zone. We presented potential geographic distributions based on ecological niche models for several species of dobsonflies. Results suggested that the geographic range of dobsonflies in the transition zone is associated to mountainous formations and that most species favor warm climates with higher precipitation rates. Climate types tend to be important for species that show narrow geographic ranges, but precipitation tends to be the most important variable to explain species dispersion. Overall, our models support the dispersion of dobsonflies from the Neotropics to North America and explain the two endemic species in Mexico as the result of the formation of the transition zone.

## Introduction

The dobsonflies are an interesting group of megalopterans belonging to the subfamily Corydalinae (Megaloptera, Corydalidae). Dobsonflies have aquatic larvae and terrestrial pupae and adults (Cover and Bogan 2015). Larvae of some species are sensitive to pollution (Maki et al. 1973; Tarter et al. 2010) and good bio-indicators of freshwater ecosystem health (Nair et al. 2015). The life cycle of most dobsonflies ranges from one to two years, though some species live up to five years (Bowles 1990; Cover and Resh 2008; Álvarez 2012; Cover and Bogan 2015). Nevertheless, adult dobsonflies have a short lifespan, only one to two weeks (Hayashi 1999; Álvarez 2012; 2014; Villagomez and Contreras-Ramos 2017), and are typically nocturnal (Cover and Resh 2008; Álvarez 2014; Álvarez et al. 2017a; 2017b; 2019).

The Neartic-Neotropical transition zone is a biogeographic region in North America at the confluence of the Nearctic and Neotropical regions and includes the mountainous provinces in the central portion of Mexico (Cuervo-Robayo et al. 2020). It has been proposed that the environmental heterogeneity of this zone may produce high rates of divergence and endemism (Espinosa et al. 2008). Its mountainous character could also limit the dispersion of Neotropical species to the north and Neartic species to the south. This zone holds 9 species of the fauna of Megaloptera (Contreras-Ramos and Rosas 2014). Though in the country 13 species of Megaloptera fauna have been recorded (Alvarez 2012; Contreras-Ramos and Rosas 2014). There are three known species of the genus *Platyneuromus* Weele, 1909 (Corydalinae) that are mostly distributed across the south, in the state of Chiapas. Three species of the genus *Chloronia* Banks, 1908 (Corydalinae) are distributed in central and south-eastern Mexico. A single species of the genus *Neohermes* Banks, 1908 (Chauliodinae) is distributed in the north of the country. A sole species of the genus *Ilyobius* Enderlein, 1910 (Sialidae) is restricted to south-eastern Mexico.

Finally, five species of the genus *Corydalus* Latreille, 1802 (Corydalinae) are distributed across the country with different geographic ranges, mostly in the center and south of the country. Recently a study of Jiang et al. (2021) showed that the biogeographic evolution of the New World Megaloptera started prior to Pangea breakup with groups affected latter by the Eurogondwanan faunal interchange, suggesting different events of vicariance and divergence, and apparently, in North America several lineage distributions are compromised by the Neartic-Neotropical transition zone. However, no studies have suggested a potential distribution hypothesis for the species of Megaloptera fauna to assess specifically the effect of this transition zone.

Aquatic insects with narrow ecological requirements, low vagility and disjunct distributions such as Megaloptera represent a valuable model for testing biogeographical hypotheses by reconstructing their distribution patterns. In this study we aim to describe a possible scenario of geographic ranges for the species of dobsonflies that occur in the Neartic-Neotropical transition zone using niche models with the data available to date, aiming to understand biogeographical patterns and to propose new insights regarding the evolution and conservation of species of the order Megaloptera.

## Materials and methods

As the Neartic-Neotropical transition zone occurs in Mexico, we focus on the species of the order Megaloptera that occur in this country. We obtained records from the data base of Mexican Megaloptera SNIB-K022 (Contreras-Ramos 2000), freely available in the Mexican National Biodiversity Information System (SNIB). This system belongs to the Mexican Commission for the Knowledge and Use of Biodiversity (CONABIO). Additionally, we used occurrence records of *Corydalus* dobsonflies provided by H.A.A. from specimens held in the entomological collection of the Autonomous University of Puebla (Álvarez 2012). All data was debugged to avoid redundancy in the modelling by selecting the records of occurrence that had a separation of three or more geographical degrees of distance between them (i.e., separation in geographical coordinates).

We used the data mentioned above in the software MaxEnt to make ecological niche models (ENM). MaxEnt is a software based on a maximum entropy method that can be used with incomplete data for prediction-making (Phillips et al. 2006). The software integrates both categorical and continuous environmental data and species occurrence data in ENM aiming to create species’ potential distribution models (Phillips et al. 2004; Phillips et al. 2006). MaxEnt was chosen here among other software because MaxEnt is fast and simple in software implementation, and it has the advantage of detail of prediction due to the continuous nature of the resulting models. Overall, whilst there may be constraints derived from data availability, it estimates the probability distribution for a species’ occurrence that is most spread out (Papes and Gaubert 2007).

We included in the models the accepted climates and ecoregions of Mexico (CONABIO), and the 19 climate variables of BIOCLIM data sets developed by Hijmans et al. (2005). BIOCLIM data sets characterize global climates using average monthly weather station data (available at different spatial resolutions). Bioclimatic variables are derived from the monthly temperature and rainfall values (biologically meaningful variables). The bioclimatic variables represent annual trends (e.g., mean annual) seasonality (e.g., annual range) and extreme or limiting environmental factors (e.g., temperature of the coldest and warmest month, and precipitation of the wet and dry quarters; being a quarter a period of three months i.e., 1/4 of the year). The codes for the 19 BIOCLIM variables are showed in Table 1.

**Table 1.** Codes and names of the 19 BIOCLIM variables (Hijmans et al. 2005).

| Code | BIOCLIM variable |
| --- | --- |
| BIO1 | Annual Mean Temperature |
| BIO2 | Mean Diurnal Range (Mean of monthly (max temp - min temp)) |
| BIO3 | Isothermality (BIO2/BIO7) ($\times 100$) |
| BIO4 | Temperature Seasonality (standard deviation $\times 100$) |
| BIO5 | Max Temperature of Warmest Month |
| BIO6 | Min Temperature of Coldest Month |
| BIO7 | Temperature Annual Range (BIO5-BIO6) |
| BIO8 | Mean Temperature of Wettest Quarter |
| BIO9 | Mean Temperature of Driest Quarter |
| BIO10 | Mean Temperature of Warmest Quarter |
| BIO11 | Mean Temperature of Coldest Quarter |
| BIO12 | Annual Precipitation |
| BIO13 | Precipitation of Wettest Month |
| BIO14 | Precipitation of Driest Month |
| BIO15 | Precipitation Seasonality (Coefficient of Variation) |
| BIO16 | Precipitation of Wettest Quarter |
| BIO17 | Precipitation of Driest Quarter |
| BIO18 | Precipitation of Warmest Quarter |
| BIO19 | Precipitation of Coldest Quarter. |

We also included three topographic variables from the project Hydro1K. These datasets have been constructed to maximize the accuracy of hydrologic parameters (www.usgs.gov) and were used in this study because they provide ready-to-use, geo-referenced hydrographic information for regional and global-scale watershed analysis and modelling.

Finally, models were made with 75% of data occurrence for standardization and 25% for validation. We consider as a potential occurrence when a model predicted 10% of the training occurrences (the minimum allowed in MaxEnt, see Papers and Gaubert 2007). The niche model results were clipped in ArcMap 9.3 using the terrestrial ecoregions from WWF to reduce the over-prediction outside the natural habitat of these insects (Olson et al. 2001) and the resulting distributions were draw above the hydro-ecoregions from WWF (Abell et al. 2008; www.feow.org).

## Results

We obtained nine significative ENM (AUC parameter upper to 0.95) for *Chloronia mexicana* Stitz, 1914, *Chloronia mirifica* Navás, 1925, *Chloronia pallida* (Davis), 1903, *Corydalus bidenticulatus* Contreras-Ramos, 1998, *Corydalus luteus* Hagen, 1861, *Corydalus magnus* Contreras-Ramos, 1998, *Corydalus peruvianus* Davis, 1903, *Corydalus texanus* Banks, 1903, and *Platyneuromus soror* (Hagen), 1861.

The predicted distributions of the species resulted primarily from environmental variables and terrestrial ecoregions, and overall, model distributions match with hydro-ecoregions. Accordingly, of the 19 BIOCLIM variables, the variables that contributed the most to models were: temperature seasonality (BIO4), mean temperature of coldest quarter (BIO11), and mean temperature of coldest month (BIO6). In addition, Table 2 summarizes the climates of Mexico that had higher incidence in the models for all species, and Table 3 shows the climates that had higher effect on the models for each species (from most to least). Overall, semi-warm subtropical climate types had the highest incidence in almost all species followed by semi-warm tropical and semi-cold subtropical.

**Table 2.**
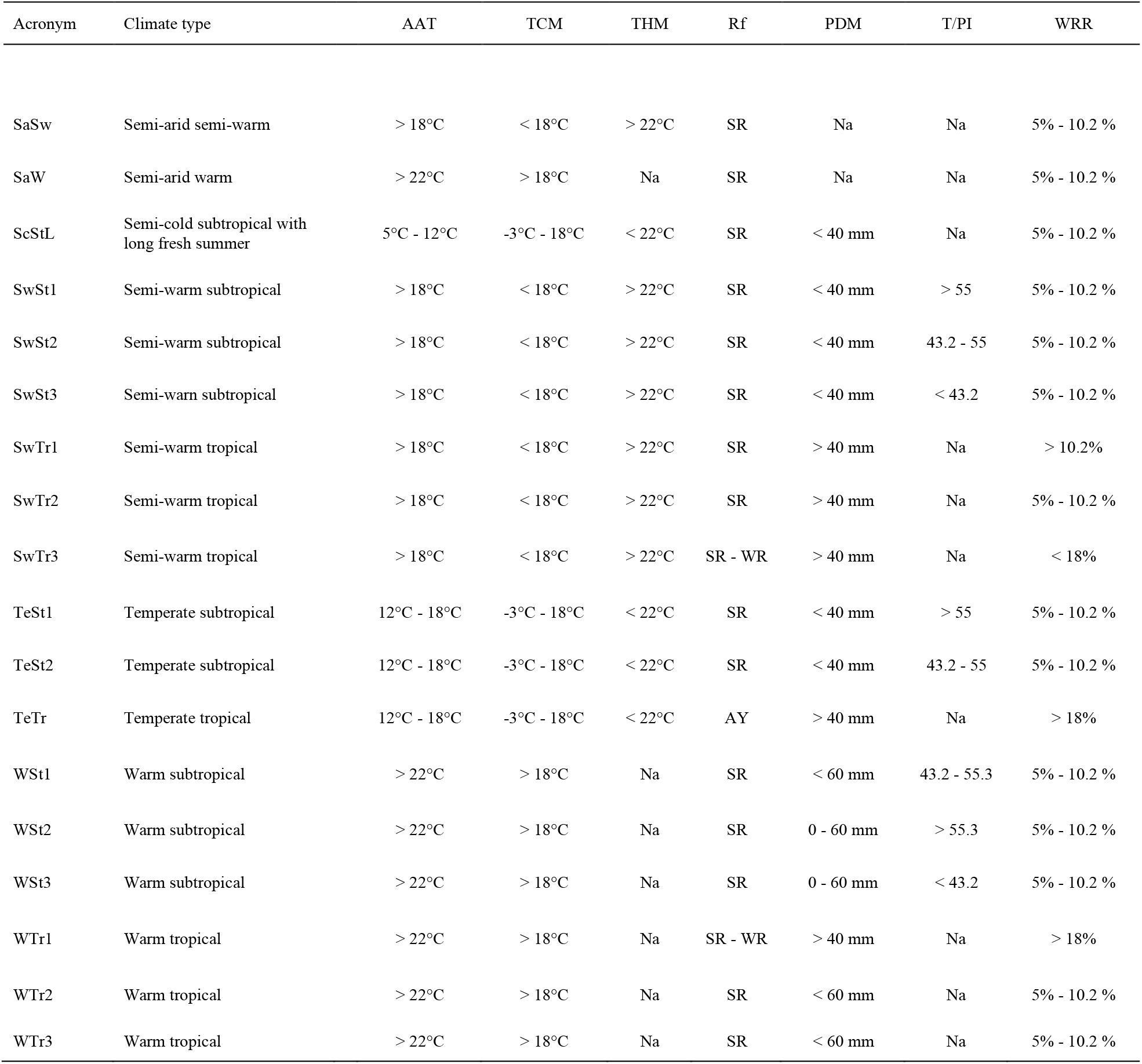
Climates of Mexico (from CONABIO) that had higher incidence in the models for all species. Climate types and their general characteristics: annual average temperature (AAT), temperature of the coldest month (TCM), temperature of the hottest month (THM), rainfall (Rf), precipitation of the driest month (PDM), temperature/precipitation index (T/PI), winter rainfall rate (WRR). Rainfall periods: summer rainfall (SR), winter rainfall (WR), all year rainfall (AY).

| Acronym | Climate type | AAT | TCM | THM | Rf | PDM | T/PI | WRR |
| --- | --- | --- | --- | --- | --- | --- | --- | --- |
| SaSw | Semi-arid semi-warm | > 18°C | < 18°C | > 22°C | SR | Na | Na | 5% - 10.2 % |
| SaW | Semi-arid warm | > 22°C | > 18°C | Na | SR | Na | Na | 5% - 10.2 % |
| ScStL | Semi-cold subtropical with long fresh summer | 5°C - 12°C | -3°C - 18°C | < 22°C | SR | < 40 mm | Na | 5% - 10.2 % |
| SwSt1 | Semi-warm subtropical | > 18°C | < 18°C | > 22°C | SR | < 40 mm | > 55 | 5% - 10.2 % |
| SwSt2 | Semi-warm subtropical | > 18°C | < 18°C | > 22°C | SR | < 40 mm | 43.2 - 55 | 5% - 10.2 % |
| SwSt3 | Semi-warm subtropical | > 18°C | < 18°C | > 22°C | SR | < 40 mm | < 43.2 | 5% - 10.2 % |
| SwTr1 | Semi-warm tropical | > 18°C | < 18°C | > 22°C | SR | > 40 mm | Na | > 10.2% |
| SwTr2 | Semi-warm tropical | > 18°C | < 18°C | > 22°C | SR | > 40 mm | Na | 5% - 10.2 % |
| SwTr3 | Semi-warm tropical | > 18°C | < 18°C | > 22°C | SR - WR | > 40 mm | Na | < 18% |
| TeSt1 | Temperate subtropical | 12°C - 18°C | -3°C - 18°C | < 22°C | SR | < 40 mm | > 55 | 5% - 10.2 % |
| TeSt2 | Temperate subtropical | 12°C - 18°C | -3°C - 18°C | < 22°C | SR | < 40 mm | 43.2 - 55 | 5% - 10.2 % |
| TeTr | Temperate tropical | 12°C - 18°C | -3°C - 18°C | < 22°C | AY | > 40 mm | Na | > 18% |
| WSt1 | Warm subtropical | > 22°C | > 18°C | Na | SR | < 60 mm | 43.2 - 55.3 | 5% - 10.2 % |
| WSt2 | Warm subtropical | > 22°C | > 18°C | Na | SR | 0 - 60 mm | > 55.3 | 5% - 10.2 % |
| WSt3 | Warm subtropical | > 22°C | > 18°C | Na | SR | 0 - 60 mm | < 43.2 | 5% - 10.2 % |
| WTr1 | Warm tropical | > 22°C | > 18°C | Na | SR - WR | > 40 mm | Na | > 18% |
| WTr2 | Warm tropical | > 22°C | > 18°C | Na | SR | < 60 mm | Na | 5% - 10.2 % |
| WTr3 | Warm tropical | > 22°C | > 18°C | Na | SR | < 60 mm | Na | 5% - 10.2 % |

**Table 3.** Climates (acronyms) with higher incidences on the niche models for each species.

| Megaloptera species | Climates |
| --- | --- |
| *Chloronia mexicana* | SwTr3, SwSt2, WSt2, SwSt1, WSt1, SwSt3 |
| *Chloronia mirifica* | SwTr3, SwTr2, WTr1, TeTr, WTr2 |
| *Chloronia pallida* | SwSt2, WSt1, SwSt1, SwSt3, WSt3 |
| *Corydalus bidenticulatus* | SwSt2, SwSt1, WSt1, SwSt3, SaW |
| *Corydalus luteus* | SwSt2, SwSt1, SwSt3, TeSt2, SwTr3, SwTr2, TeSt1 |
| *Corydalus magnus* | ScSt, SwSt1, TeSt1, TeSt2, SwSt2 |
| *Corydalus peruvianus* | ScSt, SwSt2, SwTr3, SwSt1, WSt2 |
| *Corydalus texanus* | SwSt2, SwSt1, TeSt2, SwSt3, SaSw |
| *Platyneuromus soror* | SwSt1, ScSt, TeSt1, SwSt2, SwTt2 |

## Specific distribution patterns based on modelling

### Chloronia mexicana

The species was distributed in regions with semi warm and warm climates mostly, which show high temperatures and moderate to high rainfall (Table 3). Data from niche models show that the geographic range of *C. mexicana* is associated with the Sierra Madre Oriental, the gulf coastal plain, the Balsas depression, the east section of the volcanic belt, the Sierra Madre de Oaxaca, the Tabasco plain, the south section of Yucatan platform and the north section of Chiapas highlands (Figure 1). Accordingly, this species is associated with eight hydro-ecoregions, Quintana Roo – Motagua (202), Upper Usumacinta (174), Grijalva – Usumacinta (173), Coatzacoalcos (172), Papaloapan (171), Rio Balsas (169), Panuco (167) and partly reaching Sierra Madre del Sur (170) (Figure 1, Table 4).

**Figure 1.**
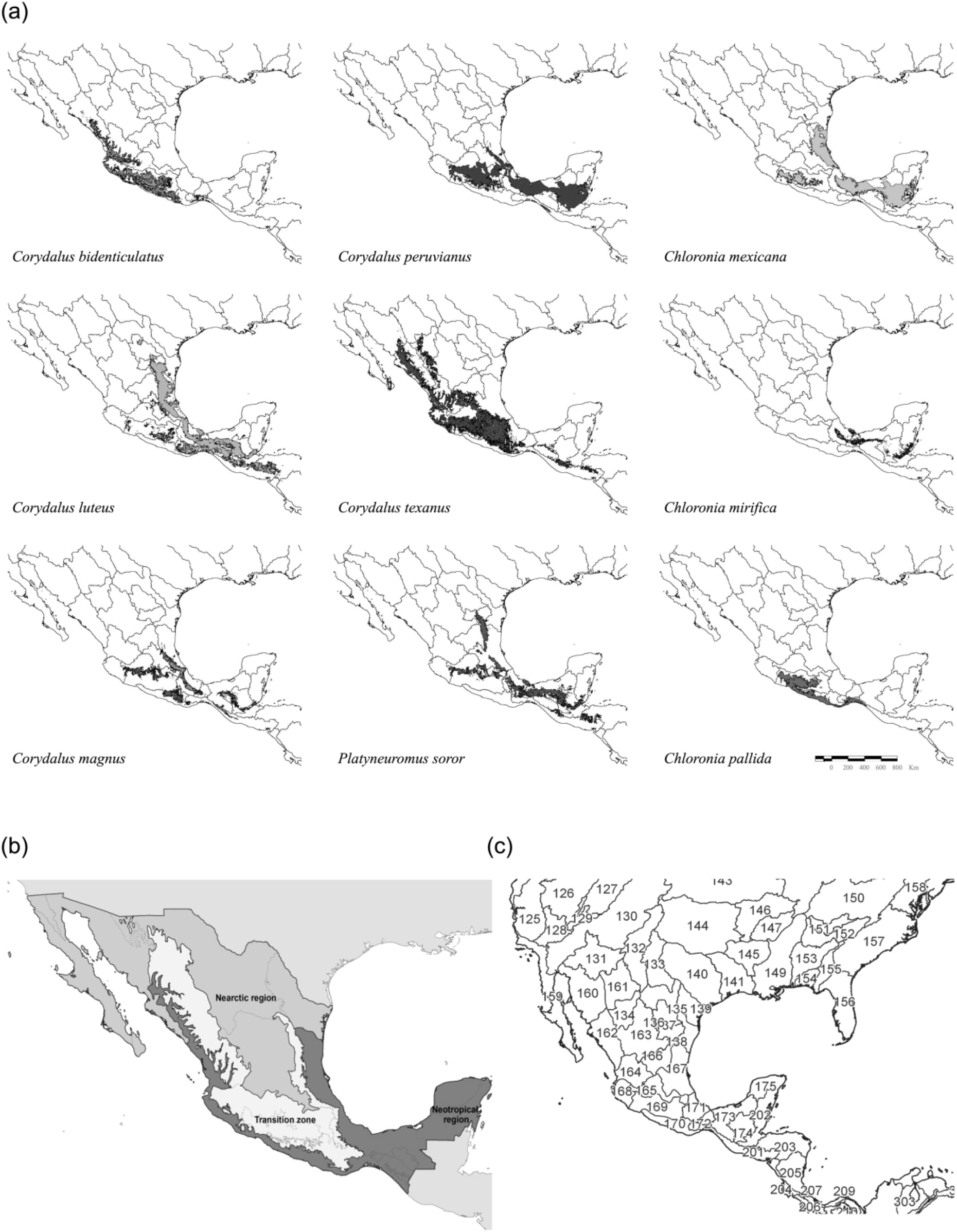
Neartic-Neotropical transition zone maps. Potential geographic distribution of *Chloronia, Corydalus*, and *Platyneuromus* species resulted from modelling (a). Biogeographic ecoregions separating the Neartic, Neotropical and the transition zone (b), and freshwater ecoregions in the area (c). Numbers represent each hydro-ecoregion’s ID number. See ID numbers for the hydro-ecoregions of Mexico in Table 4.

**Table 4.**
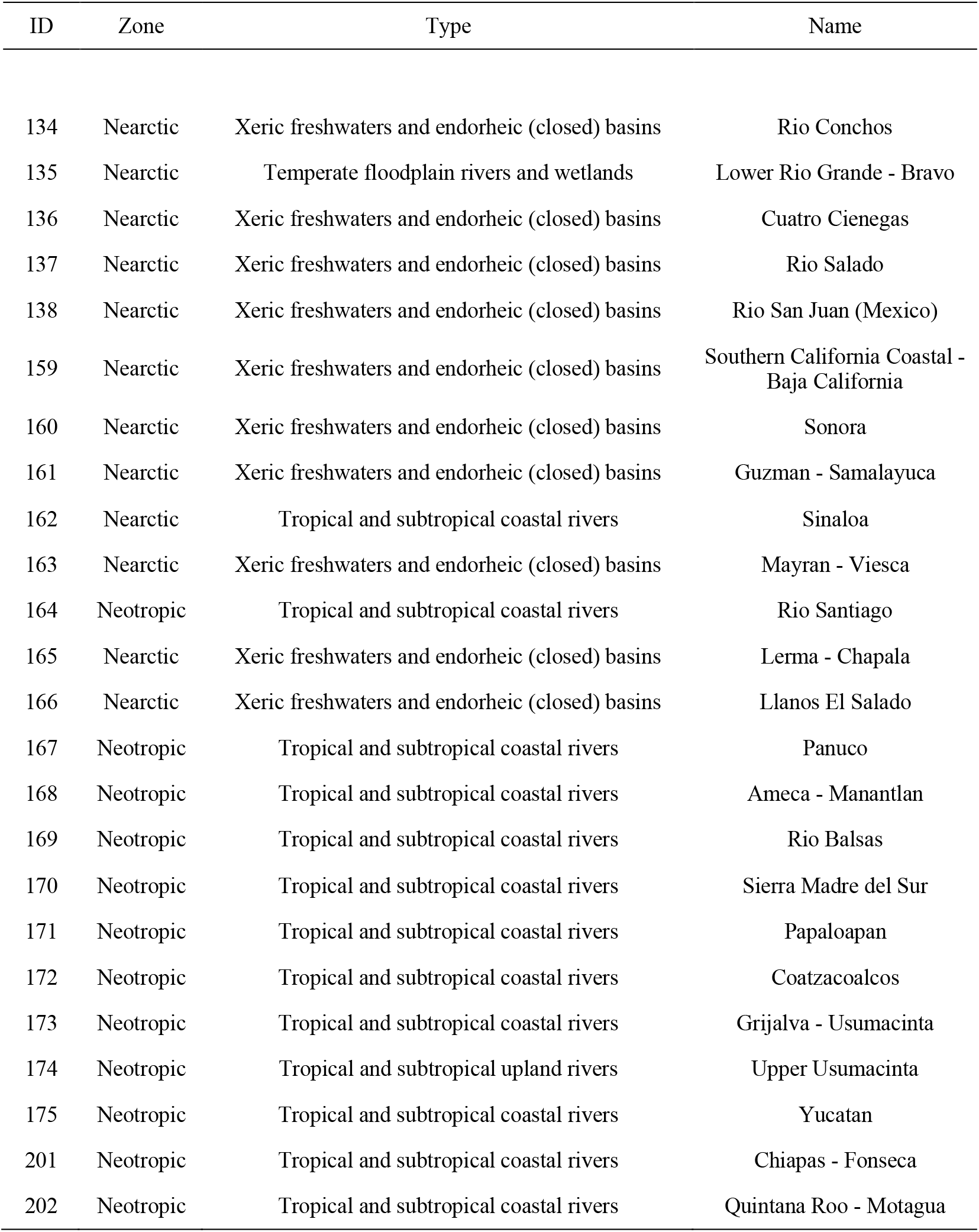
Hydro-ecoregions present in the Neartic-Neotropical transition zone. Information extracted from WWF (regions for Mexico, www.feow.org).

### Chloronia mirifica

The species was distributed in regions with tropical climates that show semi-warm to warm temperatures and high rainfall (Table 3). Data from niche models show that the geographic range of *C. mirifica* is associated with the Sierra Madre de Oaxaca, the north section of the Sierra Madre de Chiapas and the Chiapas highlands, and a restricted area in Los Tuxtlas formation (Figure 1). Accordingly, this species is associated with five hydro-ecoregions, Quintana Roo – Motagua (202), Upper Usumacinta (174), Grijalva – Usumacinta (173), Coatzacoalcos (172) and Papaloapan (171) (Figure 1, Table 4).

### Chloronia pallida

This species is endemic to Mexico where it is distributed in regions with semi warm climates, which show moderate to high temperatures and moderate rainfall (Table 3). Data from niche models show that the geographic range of *C. pallida* is associated with the south section of the Sierra Madre Oriental, the volcanic belt, the east section of the Sierra Madre del Sur, the north section of Sierra Madre de Oaxaca and the north section of the Sierra Madre de Chiapas and the Chiapas highlands (Figure 1). Accordingly, this species is associated with three hydro-ecoregions, Chiapas – Fonseca (201), Sierra Madre del Sur (170) and Rio Balsas (169) (Figure 1, Table 4).

### Corydalus bidenticulatus

This species also is endemic to Mexico where it is distributed in regions with semi-warm to arid climates, which show high temperatures and moderate to low rainfall (Table 3). Data from niche models show that the geographic range of *C. bidenticulatus* is associated with the Sierra Madre del Sur, the south section of the Sierra Madre Occidental, the pacific coastal lowlands, the volcanic belt and the Balsas depression (Figure 1). Accordingly, this species is associated with eight hydro-ecoregions, Chiapas – Fonseca (201), Sierra Madre del Sur (170), Rio Balsas (169), Ameca – Manantlan (168), Lerma – Chapala (165), Rio Santiago (164), Sinaloa (162) and partly reaching Coatzacoalcos (172) (Figure 1, Table 4).

### Corydalus luteus

The species was distributed in regions with semi-warm to temperate climates, which show moderate rainfall (Table 3). Data from niche models show that the geographic range of *C. luteus* is associated with the south section of gulf coastal plain, the Sierra Madre Oriental, the north of the Tabasco plain, part of the Balsas depression, the Chiapas highlands, the Sierra Madre de Chiapas and part of the Sierra Madre del Sur (Figure 1). Accordingly, this species is associated with thirteen hydro-ecoregions, Quintana Roo – Motagua (202), Chiapas – Fonseca (201), Upper Usumacinta (174), Grijalva – Usumacinta (173), Coatzacoalcos (172), Papaloapan (171), Sierra Madre del Sur (170), Rio Balsas (169), Panuco (167), Rio San Juan (138), Rio Salado (137), Lower Rio Grande – Bravo (135) and partly reaching Lerma – Chapala (165) (Figure 1, Table 4).

### Corydalus magnus

The species was distributed in regions with semi-cold to semi-warm climates, which show moderate low temperatures and moderate high rainfall (Table 3). Data from niche models show that the geographic range of *C. magnus* is associated with the south section of the Sierra Madre Oriental, the volcanic belt, the east section of the Sierra Madre del Sur, the north section of Sierra Madre de Oaxaca, and the north section of the Sierra Madre de Chiapas and Chiapas highlands (Figure 1). Accordingly, this species is associated with eleven hydro-ecoregions, Quintana Roo – Motagua (202), Chiapas – Fonseca (201), Upper Usumacinta (174), Grijalva – Usumacinta (173), Coatzacoalcos (172), Papaloapan (171), Sierra Madre del Sur (170), Rio Balsas (169), Ameca – Manantlan (168); Panuco (167) and Lerma – Chapala (165) (Figure 1, Table 4).

### Corydalus peruvianus

The species was distributed in regions with semi-cold to warm climates, which show moderate rainfall (Table 3). Data from niche models show that the geographic range of *C. peruvianus* is associated with the south section of the Sierra Madre Oriental and the gulf coastal plain, the volcanic belt, the mesa central, the Balsas depression, the Tabasco plain, the Chiapas highlands and the south section of Yucatan platform (Figure 1). Accordingly, this species is associated with eleven hydro-ecoregions, Quintana Roo – Motagua (202), Chiapas – Fonseca (201), Upper Usumacinta (174), Grijalva – Usumacinta (173), Coatzacoalcos (172), Papaloapan (171), Rio Balsas (169), Ameca – Manantlan (168); Panuco (167), Lerma – Chapala (165) and partly reaching Sierra Madre del Sur (170) (Figure 1, Table 4).

### Corydalus texanus

The species was distributed in regions with semi-warm to semi-arid climates, which show moderate to low rainfall (Table 3). Data from niche models show that the geographic range of *C. texanus* is associated with the Sierra Madre Occidental, the pacific coastal lowlands, the east and south section of the mesa del norte, the mesa central and the volcanic belt, the south section of the Sierra Madre Oriental, the Balsas depression, the east face of the Sierra Madre del Sur and a small section of the Sierra Madre de Chiapas (Figure 1). Accordingly, this species is associated with eighteen hydro-ecoregions, Quintana Roo – Motagua (202), Chiapas – Fonseca (201), Upper Usumacinta (174), Grijalva – Usumacinta (173), Coatzacoalcos (172), Papaloapan (171), Sierra Madre del Sur (170), Rio Balsas (169), Ameca – Manantlan (168); Panuco (167), Llanos El Salado (166), Lerma – Chapala (165), Rio Santiago (164), Mayran – Viesca (163), Sinaloa (162), Guzman – Samalayuca (161), Sonora (160) and Rio Conchos (134) (Figure 1, Table 4).

### Platyneuromus soror

The species was distributed in regions with semi-warm/semi-cold to temperate climates, which show moderate rainfall (Table 3). Data from niche models show that the geographic range of *P. soror* is associated with the west face and the south section of the Sierra Madre Oriental, the volcanic belt, and part of the mesa central, the north section of the Sierra Madre de Oaxaca, the Sierra Madre de Chiapas and the Chiapas highlands and a restricted area in Los Tuxtlas formation (Figure 1). Accordingly, this species is associated with fourteen hydro-ecoregions, Quintana Roo – Motagua (202), Chiapas – Fonseca (201), Upper Usumacinta (174), Grijalva – Usumacinta (173), Coatzacoalcos (172), Papaloapan (171), Rio Balsas (169), Ameca – Manantlan (168); Panuco (167), Llanos El Salado (166), Lerma – Chapala (165), Rio San Juan (138); Lower Rio Grande – Bravo (135) and partly reaching Sierra Madre del Sur (170) (Figure 1, Table 4).

## Discussion

Our results suggested that the geographic range of Corydalinae (dobsonflies) in the Neartic-Neotropical transition zone seems to be closely associated to mountainous formations, and that model distributions match with hydro-ecoregions (Abell et al. 2008). Accordingly, our results also show that most species favor warm climates with higher precipitation rates. Climate types tend to be important for species that show narrow geographic ranges such as *C. magnus* with a preference for cold and temperate climates or *C. mirifica* with a preference for tropical climates. However, precipitation tends to be the most important variable to explain species dispersion, i.e., species tend to disperse heading to regions with higher amounts of rainfall.

As previously suggested by Contreras-Ramos (1998) and Álvarez (2012), there is a pattern of geographical ranges for the species of Megaloptera in this zone, specifically our results suggested that almost all species are present in the Neotropical south of the country. Then, within the Neartic-Neotropical transition zone, near to the Balsas depression and the volcanic belt the geographic range of the species diverge. For example, regarding the genus *Corydalus*, three species *C. peruvianus*, *C. magnus* and *C. luteus* remain in the center, east, and south-east. Their geographic ranges are similar, broadly overlap, and share almost the same hydro-ecoregions, especially *C. peruvianus* and *C. magnus*, which prefer semi-cold climates. However, *C. magnus* had a more restricted range due to its preference for temperate climates. These three species range from the Neotropical region to the center of the transition zone and disperse within the Sierra Madre Oriental, being *C. luteus* and *C. peruvianus* the ones that reach the Gulf of Mexico. Interestingly, these three species have an origin in Central and South America (Contreras-Ramos 1998; 2011). Conversely, two species *C. texanus* and *C. bidenticulatus* remain in the south, center, and west. They do not share all the same hydro-ecoregions, in this case, *C. texanus* is the most widespread species in the country and therefore in the transition zone. In particular, *C. texanus* is distributed over the Neotropical south-west and center-west, but also over the center of the transition zone surrounding the Sierra Madre Occidental. *Corydalus bidenticulatus* appears restricted to the Neotropical west and the center of the transition zone. Regarding the genus *Chlorinia*, only *C. mexicana* disperses across the Neotropical south and east into the transition zone following the Sierra Madre Oriental and through the center of the transition zone. The rest of species remain in the Neotropical south. Finally, *Platyneuromus soror*, which is the most widespread species in that genus, shares almost the same distributional pattern and hydro-regions with *C. magnus*. Indeed, it is common to find this species sharing the same habitat and be collected in the same regions (see Álvarez et al. 2019).

In addition, the distribution of the two endemic species of *Corydalus* in Mexico can be explained by our analysis, i.e., *C. bidenticulatus* and *C. pallida*. Our results showed a relationship of their distribution pattern with the Neartic-Neotropical transition zone and their speciation may be a product of that zone. We hypothesize that an ancestral species in both *C. luteus, C. magnus, C. bidenticulatus* group (Contreras-Ramos 1998; 2011) and *C. mirifica, C. gloriosoi, C. pallida, C. mexicana* group (Penny and Flint 1982) dispersed though the area known as the Tehuantepec isthmus (hydro-ecoregions: Coatzacoalcos (172), Papaloapan (171), Sierra Madre del Sur (170), and Chiapas-Fonseca (201)), then reaching the Balsas depression. It is there at the Balsas depression where a speciation event might occur as an effect of the orogeny that gave rise to the Sierras Madre around the mid-Tertiary (as suggested by Contreras-Ramos 1998). Then the new species dispersed, i.e., *C. bidenticulatus* to the north and south-east and *C. pallida* to the south and east, both following the Neotropical Pacific coast and the Balsas depression.

Overall, it is possible that while precipitation is a crucial, if not most important, factor influencing the current geographic distribution of these various species, other potential influences might include the degree of habitat conservation (pollution and fragmentation) and the continuum of the total ecosystem (Estrada and Coates-Estrada 1996; Cayuela et al. 2006; Arriaga-Weiss et al. 2008). For example, it has been shown that within landscapes that are less homogeneous, mountainous formations are not extremely modified by human activities. These rugged landscapes naturally provide zones where species can inhabit with lower perturbation compared to other types of regions, in fact, some conservational approaches have been successful on this type of landscapes and corridors (DeClerck et al. 2010). This could explain the tendencies in the geographic ranges of Megaloptera fauna within the Neartic-Neotropical transition zone. Though, the Sierra Madre Occidental, the Sierra Madre Oriental, the volcanic belt, the Sierra Madre del Sur and their sub-divisional lower formations, provide a continuous of aquatic habitats that megalopterans may inhabit (Álvarez 2012), but specific climate types might constrain the distribution of the species for lower (near sea level) or higher altitudes. So, if the species studied here are too sensitive to these factors, then the real geographic range of Corydalinae species could be minor than the model prediction in our analysis.

Here we provide the first approach to understand biogeographic distribution of the Megaloptera from the Neartic-Neotropical transition zone. However, more efforts are needed to achieve biogeographical modelling of the entire fauna of Megaloptera in this region of the Americas. Field data and large-scale approaches such as continental scale will allow a better understanding of the biogeographic patterns and evolution of this insect order. Nonetheless, our results are in agreement with the biogeographical hypothesis for New World Megaloptera proposed by Contreras-Ramos (1998) and Penny (1993), and the results provided by Jiang et al. (2021), which showed that the biogeographical patterns in North America are the result of two main events, firstly, Sialidae, Corydalinae, and Chauliodinae originated previous to the breakup of Pangea (late Jurassic ~160-145 Ma). Chauliodinae and Sialidae dispersed through Laurentia and West Gondwana and then dispersed across the area until the new continent was formed, with subsequent events of vicariance. Conversely, Corydalinae and especially the *Corydalus* lineage dispersed from Asia via Europe and Africa to South America as a result of the second Eurogondwanan connection (Late Cretaceous ~100-90 Ma) (Oliveira et al. 2010; Jiang et al. 2021). Secondly, they dispersed through South America then to Central and North America, with subsequent events of vicariance, which for North America was the rise and formation of the Neartic-Neotropical transition zone.

## Acknowledgments

The authors want to thank Manolo Tierno de Figueroa, Lorelí Carranza, and an anonymous reviewer for their helpful comments on an earlier version of the manuscript. H.A.A. thanks Gemma Clemente Orta for her help and comments on figures.

